# Is there a ubiquitous spectrolaminar motif of local field potential power across primate neocortex?

**DOI:** 10.1101/2024.09.18.613490

**Authors:** C. A. Mackey, K. Duecker, S. Neymotin, S. Dura-Bernal, S. Haegens, A. Barczak, M. N. O’Connell, S.R. Jones, M. Ding, A.S. Ghuman, C.E. Schroeder

## Abstract

Mendoza-Halliday, Major et al.^1^ (referred to as “Mendoza-Halliday et al.” for brevity), advocate for a local field potential (LFP)-based approach to functional identification of cortical layers during “laminar” (simultaneous recordings from all cortical layers) multielectrode recordings in nonhuman primates (NHPs). They describe a “ubiquitous spectrolaminar motif” in the primate neocortex: 1) 75-150 Hz power peaks in the supragranular layers, 2) 10-19 Hz power peaks in the infragranular layers and 3) the crossing point of their laminar power gradients identifies Layer 4 (L4). Identification of L4 is critical in general, but especially for *Mendoza-Halliday et al*. as the “motif” discovery is couched within a framework whose central hypothesis is that gamma activity originates in the supragranular layers and reflects feedforward activity, while alpha-beta activity originates in the infragranular layers and reflects feedback activity. In an impressive scientific effort, *Mendoza-Halliday et al*. analyzed laminar data from 14 cortical areas in 2 prior macaque studies and compared them to marmoset, mouse, and human data to further bolster the canonical nature of the motif. Identification of such canonical principles of brain operation is clearly a topic of broad scientific interest. Similarly, a reliable online method for L4 identification would be of broad scientific value for the rapidly increasing use of laminar recordings using numerous evolving technologies. Despite *Mendoza-Halliday et al*.’s paper’s strengths, and its potential for scientific impact, a series of concerns that are fundamental to the analysis and interpretation of laminar activity profile data in general, and local field potential (LFP) signals in particular, led us to question its conclusions. Here, we address four key questions: **Q1**) Is the spectrolaminar motif ubiquitous, i.e. “found everywhere” or “universal”? **Q2**) Do features of the motif reliably identify Layer (L) 4? **Q3**) Are *Mendoza-Halliday et al.’s* newly introduced methods (FLIP and vFLIP) reliable? And **Q4**) Are the proposed biophysical mechanisms underlying the motif well justified? We used new sets of data comprised of stimulus-evoked laminar response profiles from primary and higher-order auditory cortices (A1 and belt cortex), and primary visual cortex (V1) to test these questions. The rationale for using these areas as a test bed for new methods is that, in contrast to higher-order cortical areas, their laminar anatomy and physiology have already been extensively characterized by prior studies, and there is general agreement across laboratories on key matters like L4 identification. In short, we find that *Mendoza-Halliday et al.’*s findings do not generalize well to these cortical areas. Specifically, regarding **Q1**: Though we can find a spectrolaminar gradient that is *qualitatively* consistent with *Mendoza-Halliday et al*., it is quantifiable in only 61-64% of our recordings, indicating that the motif is common but by no means universal (see “**Evaluation of the FLIP method** […]” and “**Evaluation of vFLIP** […]”). Regarding **Q2**: The motif’s high/low frequency gradient cross point identified L4 in only 29-33% of our recordings. Regarding **Q3:** FLIP and vFLIP exhibit marked variability across studies, across brain areas, and spuriously detect cortical layer inversions (see “**FLIP’s fitting process** […]” and “**Evaluation of vFLIP** […]”). Regarding **Q4:** the biophysical modeling findings cited to support *Mendoza-Halliday’s* conclusions (see “**Going forward – *in silico*** […]”) do not reproduce the LFP data trends. While our findings are in many respects at odds with those of *Mendoza-Halliday et al*., the paper already has, and will continue to spark debate and further experimentation. Hopefully this countervailing presentation will lead to robust collegial efforts to define optimal strategies for applying laminar recording methods in future studies.

## Evaluation of the FLIP method in new data sets

We applied *Mendoza-Halliday et al*.’s main analytic strategy, **F**requency-based **L**ayer **I**dentification **P**rocedure (FLIP), to three data sets recorded from primary (A1) and belt (secondary) auditory cortex (2 monkeys) and V1 (2 monkeys). These areas were chosen because, unlike higher-order areas such as PFC and LIP: 1) their laminar field potential, current source density (CSD) and multiunit activity (MUA) profiles have been thoroughly characterized by a number of labs, 2) they exhibit robust sensory-evoked responses with laminar activation patterns that reliably identify the positions of the supragranular, granular and infragranular layers ^2–6^ and 3) there is general agreement across labs on this point ^3,7–9^. The prior work in these areas, which incorporates interlocking physiology and anatomical reconstruction, provides substantial ground truth for evaluating Mendoza-Halliday et al.’s methods and conclusions. Positive findings here would not guarantee generalization to all cortical areas, but negative findings would argue against generality.

*Mendoza-Halliday et al*. contend that the crossing point between high and low frequency relative power gradients marks the location of cortical Layer 4 (L4). To identify this location, FLIP uses a linear regression approach, iterating over combinations of recording probe channels’ signals. The laminar region with the highest goodness-of-fit value (G) for low vs high frequency power regression lines with slopes of opposite signs determines the range of probe channels to be used as the “regression region” within which to search for the location where the power gradients cross. We encountered several problems with this approach.

### First, *FLIP does not reliably identify L4*

**Figures 1A-C** show findings from a representative site in A1 where current source density (CSD) and multiunit activity (MUA) profiles were used to identify the position of L4 (white lines), for comparison with FLIP-defined layer locations (**Fig 1A**). Based on laminar distributions of relative LFP power in the 75-150Hz and 10-19 Hz ranges for this site, time-locked to the stimulus and normalized to the channel with maximal power, the FLIP-defined laminar power gradient crossing falls outside of (in this case below) L4 (**Fig 1B**). Quantification of FLIP-defined L4 locations for the subset of profiles it is able to fit (**Fig 1C**), registered in the framework of the CSD/MUA-defined layer scheme at each site for our A1, auditory belt and V1 data sets (**Fig 1D**), shows that this finding is general across areas: for cases in which the FLIP algorithm could find a fit (64% overall), the power gradient crossing falls in L4 in only 29% of the cases overall (32% in A1, 40% in belt and 8% in V1). In the majority of cases (71%), the FLIP-defined power gradient crossing point falls in the infragranular layers. Critical here is that this finding depended on examination of individual penetrations/experiments, rather than aligning and averaging data across experiments and brain areas (a technique utilized by *Mendoza-Halliday et al*.).

**Fig 1.**
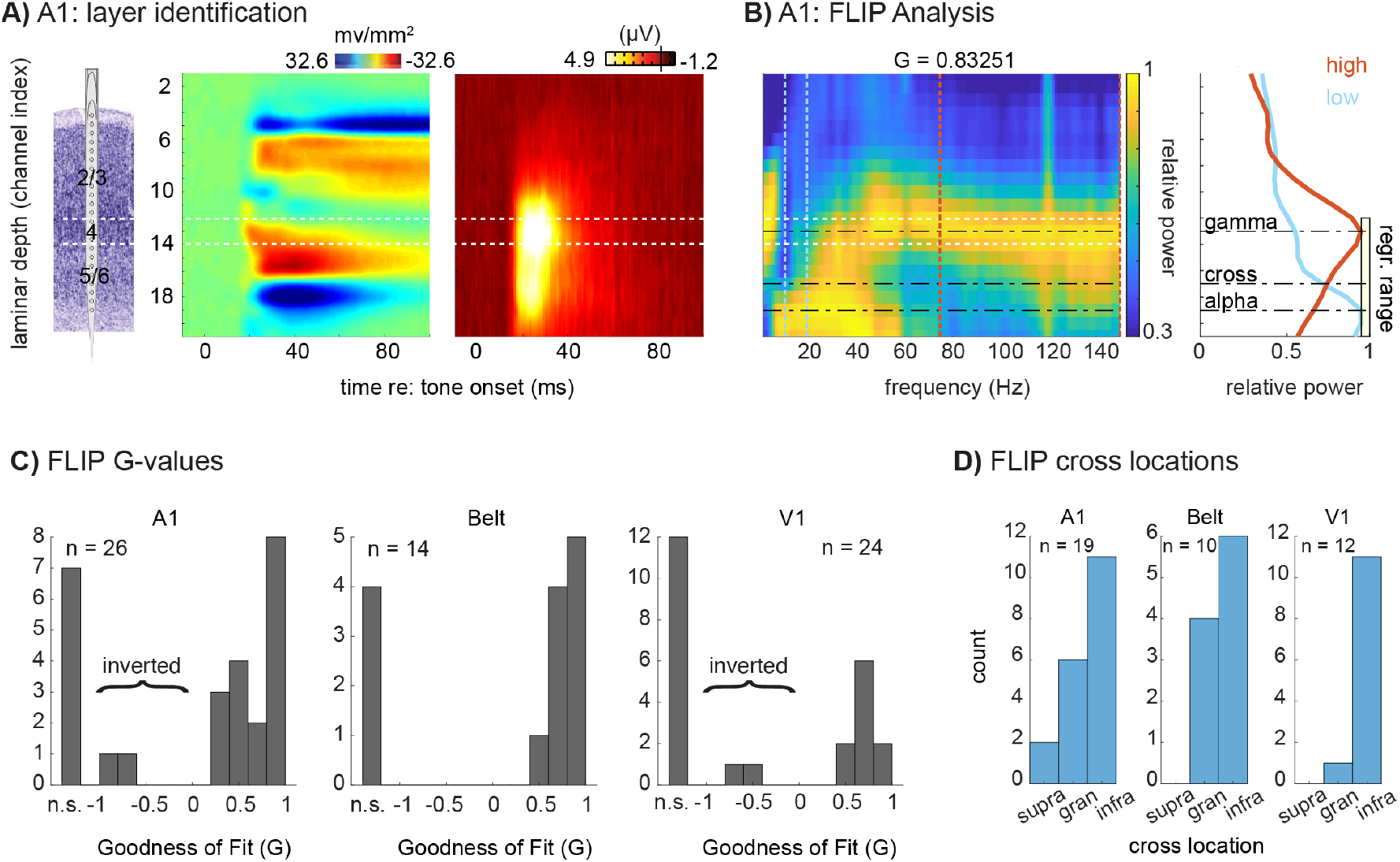
Frequency-based Layer Identification Procedure (FLIP) analysis in auditory and visual cortices. **A)** Representative tone-evoked CSD and MUA profiles used to identify cortical layers in A1; Layer 4 is outlined in white. Laminar identification used methods established by prior studies. Specifically, the electrode array was positioned to straddle the region exhibiting significant (non-overlapping trial-wise confidence intervals between the pre- and post-stimulus periods) CSD activation. Layer 4’s location was defined as the channel in the middle third portion of the active region with the earliest current sink onset, within 100 microns (+/-1 channel in this case) of the peak amplitude MUA site. **B)** Relative LFP power at each frequency, epoched -300 to 300 ms around stimulus onset, displayed as a color plot (left) with the associated goodness of fit (*G*) value generated by the FLIP algorithm; line plots (right) depict the laminar power gradients for alpha-beta (blue line, 10-20 Hz) and broadband gamma (red line; 75 - 150 Hz). The dashed lines indicate low and high frequency peaks picked by FLIP, and the power gradient crossing point. The yellow box indicates the FLIP-selected range of channels over which it conducts the regression. **C)** Goodness of fit (*G*-values) resulting from FLIP for A1, Belt, and V1. Values could be not significant (“n.s.”), negative (inverted motif), or positive. **D)** FLIP-defined power gradient crossing locations for the subset of sites with significant fits by FLIP, for A1, Belt cortex, and V1, binned into CSD/MUA-defined supragranular, granular, or infragranular compartments.

### Second, *FLIP’s fitting process is variable and error prone*

While FLIP was able to find an adequate ‘fit’ for 64% of sites overall (comparable to *Mendoza-Halliday et al*.*’s* findings), the values were highly variable across areas, with 73% of A1 sites and 71% of auditory belt sites, but only 50% of V1 sites (**Fig 1C)**. Note that a similar degree of interareal variability, and marked inter-study variability, was also evident in the shared dataset accompanying *Mendoza-Halliday et al*. (**Table S-1**). Note also, that in our data sets, FLIP finds 2 cases in A1 and two cases V1 where the G value is negative, implying that the laminar array penetrated cortex from the white matter surface, producing an “inverted” motif (**Fig 1C)** as reported by *Mendoza-Halliday et al*. Such inverted penetration is possible for peripheral V1 representations lying in the inner folds of the Calcarine Sulcus, and it produces clearly recognizable inversions of LFP, CSD and MUA profiles (see ref ^3^ Fig 4). However, this scenario is anatomically implausible for our data sets, since our A1 and V1 samples came entirely from areas lying on flat, unfolded cortical planes, and electrode penetrations were all made through the pial surface of the cortex. Observation of these “motif inversion errors” raises concern about interpretation of motif inversions in *Mendoza-Halliday et al*. (**Table S-1**).

### Consideration of the vFLIP method

The second quantitative algorithm used by *Mendoza-Halliday et al*. is “vFLIP,” a frequency-variable version of FLIP. vFLIP first runs the FLIP process over a large space of potential frequency bands and picks the best fit bands instead of using preset frequency bands the way FLIP does. ***This approach produces a multiple comparison problem and the threshold used for vFLIP must be adjusted accordingly***. *Mendoza-Halliday et al*. does not specify the threshold used for vFLIP, but it states that vFLIP identifies 81% of penetrations showing “opposing gradients of 10-19 Hz and 75-150 Hz power” across laminae, while FLIP identifies 64%. The determination of an appropriate threshold is critical, because vFLIP can find relatively high goodness of fit (G) values for random noise (**Fig 2A**). We analyzed data from noise simulations using vFLIP to determine the “G” threshold equivalent to the .265 threshold used for FLIP by *Mendoza-Halliday et al*. (**Fig 2B**); this equivalent threshold corresponds to a single penetration p-value of approximately p<.03 for noise simulations and a false discovery rate of approximately .05 in the public dataset by *Mendoza-Halliday et al*. (see **Supplement**; **1**.). When we applied this appropriate threshold, ***vFLIP identifies approximately 3% fewer penetrations than FLIP does*** (rather than 17% more) in that shared dataset. Notably, ***vFLIP findings agree with FLIP findings in only 49% of cases*** in the shared dataset (i.e., the data from figures 1-6 of *Mendoza-Halliday et al*.). “49%” refers specifically to: 1) agreement between FLIP and vFLIP on whether a penetration is “fit” (identified as having opposing gradients), 2) when both methods identify a particular penetration as fit, they specify the same (upright/inverted) orientation and 3) the electrode channels identified as the crossover by the two methods are within 200 microns of one another; this “agreement” becomes only 56% if the criterion is relaxed to 500 microns. Allowance for some variation in peak alpha-beta and gamma frequencies is a key strength of vFLIP. However, a significant factor in the divergence of FLIP and vFLIP results for the same data set is that vFLIP tends to find very narrow bands (the vast majority of bands are at the 10 Hz minimum allowed by the code) for both alpha-beta and gamma (**Fig S-1**); this also has negative consequences for vFLIP’s wider use in the study of neuronal dynamics.

**Figure 2.**
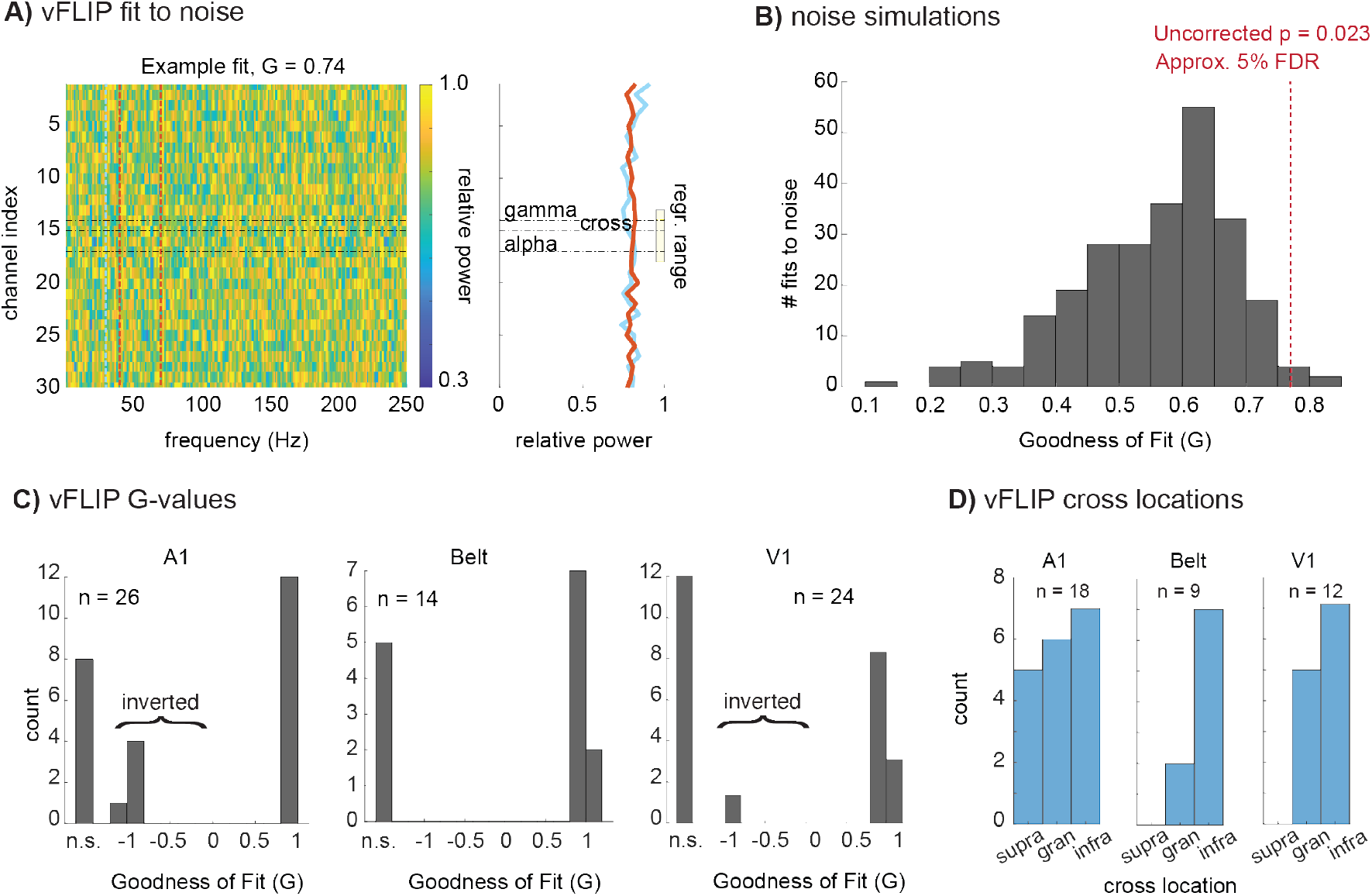
Variable Frequency-based Layer Identification Procedure (vFLIP) analysis. **A)** Representative relative power map of 1/f noise samples (30 channels, 1000 time points, 75 trials) displayed as a color plot (left) with the associated goodness of fit (*G*) value from vFLIP, 0.74. **B)** A histogram depicts the *G*-values produced by vFLIP when fit to 1/f noise 250 times. A red line depicts the statistical threshold used to attain a 5% False-discovery rate, which was then applied to the vFLIP results in auditory and visual cortices. **C)** as in Fig. 1, vFLIP cross locations determined by co-registration with CSD and MUA profiles for areas A1, Belt auditory cortex, and V1, and binned into supragranular, granular, or infragranular compartments. **D)** as in Fig. 1, *G*-values for A1, Belt, and V1, which could be not significant (“n.s.”), negative (inverted motif), or positive.

### Evaluation of vFLIP in new data sets

Application of vFLIP to our data sets re-iterates the critical problems seen with FLIP. <u>First</u>, like FLIP, ***vFLIP’s identification of Layer 4 is unreliable***. For the 61% of penetrations overall that vFLIP is able to fit (**Fig 2C**), it correctly identifies Layer 4 in well under 1/2 of the cases in A1 (33%), auditory belt (22%) and V1 (42%), (**Fig 2D**). <u>Second</u>, ***vFLIP incorrectly labels penetrations as “inverted”*** (**Fig 2C**). While FLIP was the primary analyses of the non-human primate data by *Mendoza-Halliday et al*., concerns about vFLIP are significant, as vFLIP was the primary method used for the interspecies generalization analyses.

### Broader concerns about *Mendoza-Halliday, Major et al*.’s approach

The specific analytic problems noted above connect to broader concerns about *Mendoza-Halliday et al*.’s approach and its conclusions. We address 3 concerns here and defer others to the **supplement** (**4.1-4.5**).

A <u>first</u> critical concern is establishment of ground truth in cortical layer identification. As noted above, we used a prior, broadly supported approach to laminar identification for the three low level cortical areas we used as a test bed for the FLIP and vFLIP methods. *Mendoza-Halliday et al*. proposed to link their physiological biomarker for Layer 4 (the depth of high/low frequency gradient crossing point) directly to anatomy via lesion/anatomical reconstruction. In principle this is an excellent approach, but the actual results are highly problematic (see **supplement 4.4**)

<u>A second concern is that</u> *Mendoza-Halliday et al*. used an image similarity metric to support the idea that results of vFLIP are equivalent both within and across species, however, this metric is meant to compare similarity of images based on human perceptual biases and is highly vulnerable to artifacts when it is used for comparing spectrolaminar motifs. Effects of systematically increasing the degree of blur in a pair of images on the image similarity metric used by *Mendoza-Halliday et al*. suggests that values found in Figure 8 of *Mendoza-Halliday et al*. are well within a pure noise range (**Fig S-2**).

A <u>third</u> concern is that both FLIP and vFLIP analyze LFPs recorded against a distant reference, a practice that is relatively imprecise in its spatial localization of signal origins, and thus, at odds with the objective of precisely locating L4 and differentiating activity in superficial versus deep layers of cortex^9–11^. LFP quantification measures like FLIP and the image similarity metric used by *Mendoza-Halliday et al*. produce increasing false positive rates as the spatial resolution of the measure decreases. While the threshold (*G* = 0.265) used by *Mendoza-Halliday et al*. is appropriate when spatial resolution is high, it is not adjusted for the lower spatial resolution of LFP measured against a distant reference. In the latter case the risk of false positive outcomes increases significantly (**Fig S-3**), leaving unclear the real, quantifiable percentage of laminar motifs that have the pattern referred to as “ubiquitous.” Note that the false positive rate is reduced with bipolar and CSD derivations. Studies that localize field potentials using the more spatially precise bipolar and CSD derivations ^9,11,12^ outline a paradox: alpha/beta LFP power peaks in the infragranular layers, but the strongest generators (transmembrane currents that set-up the field potential distribution in the extracellular medium) are in the supragranular layers (**Fig S-4;** see also *Mendoza-Halliday et al*.’s Supplemental Fig 9). Volume conduction effects with a distant reference pose the risk that LFP measures can mis-localize neural activity not only across cortical layers, but also across cortical areas^13,14^. Despite this, *Mendoza-Halliday et al*. refer to “layer-specific neuronal oscillations” based on their distantly referenced LFP recordings. A search for a functional method that reliably identifies specific cortical layers, and that generalizes across areas should start with methods having the best available signal localization qualities (i.e., the highest spatial resolution).

### Going forward – *in-silico* experiments to inform and extend *in-vivo* findings

*Mendoza-Halliday et al*. also discuss Neuronal Mass Modeling in an attempt to resolve the paradox between unipolar FP analysis and bipolar/CSD derivations^15^. However, the model does not seem to reproduce *Mendoza-Halliday et al*.’s experimental data well. While the increasing relative LFP power gradient for the low-frequencies from superficial to deep was somewhat approximated, the model does not reproduce the high-frequency component of the LFP, and instead estimates the location of its generator in the granular instead of the supragranular layers (this is noted by the authors^15^). Furthermore, the model CSD shows a peak in the granular layers for both the slow and fast frequencies. This is in conflict with the CSD measure derived from the data, which shows a decreasing gradient from supragranular to deep for both the high and low frequencies. Since the model does not reproduce the experimental LFP/CSD patterns, the alternative to a volume conduction interpretation, proposed by *Mendoza-Halliday et al*., lacks support. These discrepancies may be due to limitations in the design of the model, for instance, its sparse neurobiological detail, which can be avoided by using publicly available, detailed biophysical models (see **Supplement** ).

## Author Contributions

AB, MNO and CES contributed the auditory and visual cortex data sets. CAM, KD and ASG conducted primary data analyses. CAM, KD, CES, ASG, SN, and SD-B drafted the manuscript. All authors significantly contributed to the conceptualization and editing of the manuscript.

## Acknowledgments

R01DC012947, R01DC019979, P50MH109429

NIH R01NS128924-01

NIH R01MH134118-01

ARL Cooperative Agreement W911NF2220143

The authors thank Drs. Robert T. Knight, Christopher Honey and Yoshinao Kajikawa for critical comments on an earlier version of the manuscript. We also wish to acknowledge Drs. Henry Evrard, Arnaud Falchier, and John Smiley for helpful comments and advice on histological methodology.

## Data and code availability

Data and code by Mendoza-Halliday et al. are available here: https://datadryad.org/stash/dataset/doi:10.5061/dryad.9w0vt4bnp

All other code and data have been provided by the Editor.

## Ethical statement

All data were collected using procedures approved by the Nathan S. Kline Institute Institutional Animal Care and Use Committee, and followed the guidelines set forth by the NKI IACUC and USDA.

Arising from: Mendoza-Halliday, Major et al. *Nature Neuroscience* https://doi.org/10.1038/s41593-023-01554-7 (2024).

This **supplement** provides additional, in-depth explanation of points raised in the main text, as well as a more complete set of references.

## 1. vFLIP does not find more sites with a significant fit than FLIP when an appropriate threshold is used

*Mendoza-Halliday, Major et al*. ^1^ (“*Mendoza-Halliday et al*.*”* for brevity) do not state what threshold was used for vFLIP (the only threshold noted is .265 for FLIP), thus we sought to estimate an appropriate threshold for vFLIP to examine the performance of this algorithm. Because vFLIP runs FLIP many times iterating over different possible frequency ranges, there are a large number of multiple comparisons that must be taken into account when determining the threshold for vFLIP. We estimated the threshold for vFLIP that was statistically equivalent to the .265 used for FLIP to appropriately compare the performance of vFLIP to FLIP. Accordingly, we first ran FLIP 10,000 times on randomly generated pink noise (10,000 simulations of 30 channels with 1000 timepoints and 75 trials of 1/f noise). We then passed this noise through the code shared by Mendoza-Halliday et al., which first uses Fieldtrip™ to generate the spectrolaminar pattern for this noise, then uses FLIP to determine if there was the “X” shaped pattern across channels for low and high frequency power. Across the 10,000 simulations, the .265 threshold of FLIP corresponded to a false positive rate (detection of the “X” pattern in noise) of approximately 2.3%; this corresponded to an estimated false discovery rate of less than 5% in the dataset shared by Mendoza-Halliday et al.

To determine the threshold for vFLIP, we repeated this same estimation procedure 500 times (fewer simulations were used for vFLIP because vFLIP is computationally intensive and slow to run). The vFLIP threshold that gave the same false positive rate as FLIP was .76. Using this threshold on the dataset shared by Mendoza-Halliday et al., vFLIP detects 3% fewer penetrations as having the “X” pattern than FLIP, rather than 17% more (as reported by Mendoza-Halliday et al.). Therefore, the threshold used in *The Paper* was one that likely had a high false positive rate (their Fig. 2B). Similar results were seen for FLIP and vFLIP in the new dataset reported in this *Matters Arising* work, where 64% of penetrations were detected by FLIP at a .265 threshold and 61% were detected by vFLIP at a .76 threshold.

## 2. vFLIP tends towards very narrow frequency range selections in an overly broad parameter space

<u>First,</u> vFLIP’s relatively unrestricted search of the frequency band space usually results in selection of narrow (∼10 Hz) frequency bands, indicating that vFLIP operates in a parameter space with numerous potential peaks and power gradient crossing points, many of which are likely spurious (see main text **Fig. 2**).

<u>Second,</u> while allowance for some variation in peak alpha-beta and gamma frequencies is a strength of vFLIP analysis, *“low” frequencies are allowed vary from 1-70 Hz, and “high” frequencies can vary from 30-150 Hz*, resulting in frequency selections that often deviate from an alpha/beta-gamma motif (**Fig S-1**).

**Fig S-1.**
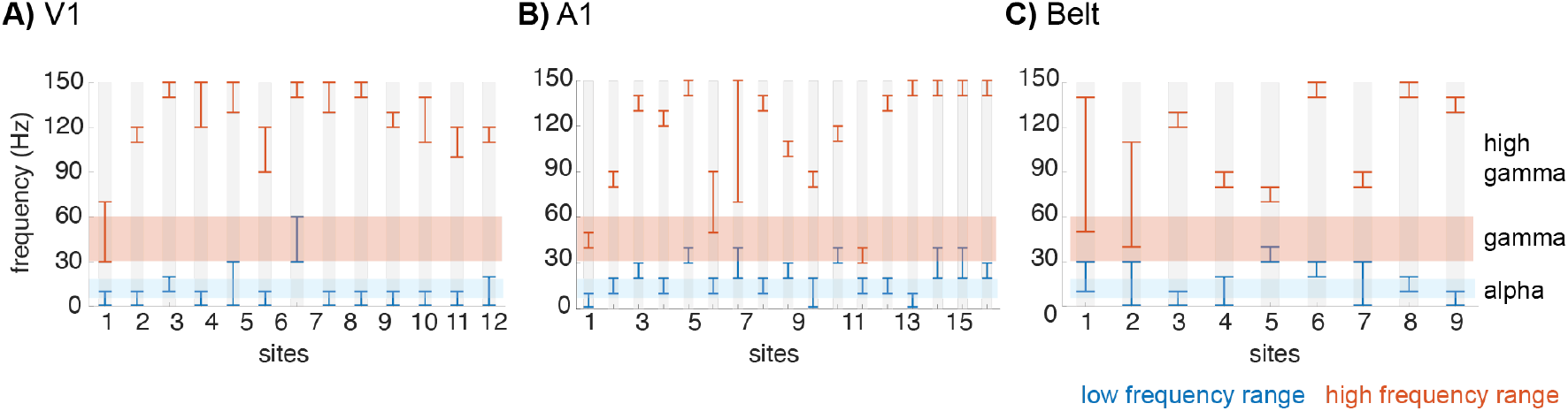
Automatically selected frequency bands from the vFLIP algorithm for each site in **A)** V1, **B)** A1, and **C)** Belt. “High” frequencies are orange, “low” frequencies are blue, and alpha/beta, gamma, and high gamma ranges are color-coded accordingly.

Notably, there are numerous cases where the “low” frequencies extend into the gamma band, and high frequencies extend well beyond the conventional gamma band of 30-60 Hz and into the 120-150 Hz high gamma band, that is likely representative of high frequency transient (e.g., spiking) rather than oscillatory activity. In fact, ***there are only a few cases in which the low/high frequencies are in the alpha-beta and gamma ranges respectively***, contrary to *Mendoza-Halliday et al*.*’s* suggestion. Because vFLIP labels any “low” frequency as “alpha/beta.” As the authors don’t report what frequencies vFLIP selected, it’s unclear if the alpha/beta-gamma motif reported in the paper is actually gamma-high gamma, delta-gamma, etc.

## 3. False positive rate of spatially imprecise LFP measures for FLIP and image similarity analysis

Key measures used by *Mendoza-Halliday et al*. produce increasing false positive rates as the spatial resolution of the measure decreases. Critically, the thresholds used in *The Paper* are only appropriate when spatial resolution is high and are not adjusted for the lower spatial resolution of LFP measures using a distant reference. This leaves unclear what is the true, quantifiable percentage of laminar motifs that have the X-like pattern that the authors call “ubiquitous.”

The SSIM image similarity metric used by *Mendoza-Halliday et al*. to establish the similarity of spectrolaminar motifs suffers from reduced specificity with increased spatial blur. **Fig S-2A** shows the image similarity using SSIM for random noise with increasing blur. The image similarity rapidly increases even for random noise as the image is blurred more. **Fig S-2B** shows an example of two spectrolaminar plots taken from *Mendoza-Halliday et al*.*’s* dataset that, *even with opposite motifs, achieve an image similarity of* .*45, higher than the reported similarity for any of the within or across species comparisons in their Figure 8i*.

**Fig S-2.**
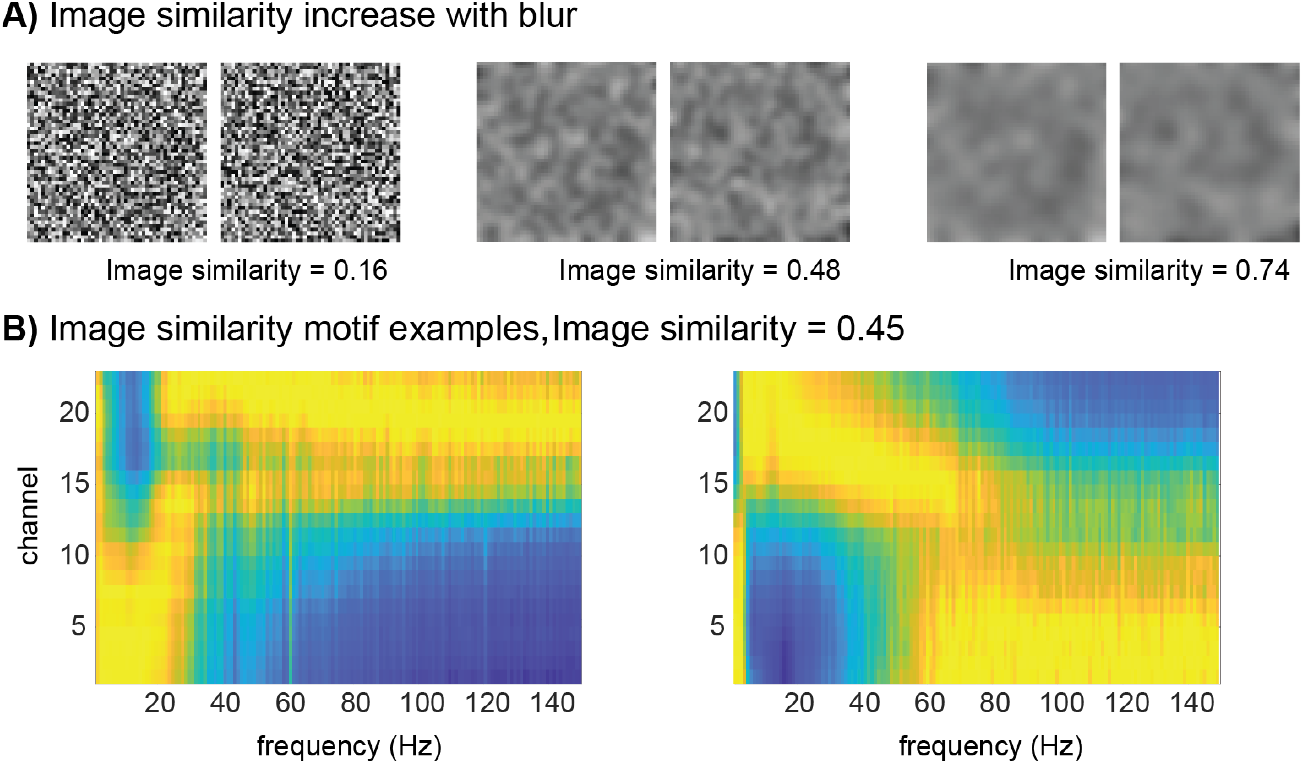
Interpretability of the Image Similarity measure. **A)** Image similarity increases for the same examples of noisy images with increasing blur. Using SSIM the similarity between random noise images was calculated with increasing blur. The image similarity values below the examples in this panel reflect the average image similarity of 10000 pairs of random images unblurred and then blurred at 2 different amounts. **B)** Image similarity for relative power in two examples probes from the publicly available datasets from *The Paper*. These two probes were chosen as having strong positive and negative G. Despite the notable differences and near inverted profiles, the image similarity reaches a value of 0.45, which is higher than the similarity for within and across species comparisons reported in *The Paper* (figure 8i).

This is not a flaw in the SSIM method, rather the SSIM method is an inappropriate measure for determining if two spectrolaminar patterns match. SSIM was developed to evaluate whether a compressed picture would be judged similarly to the original by a human observer. It thus examines factors like local contrast, luminance, and structure and incorporates Weber’s law^2^. Therefore, for example, noise with greater blur does look more similar by eye even if the underlying data are not similar at all. For assessing the similarity of spectrolaminar patterns, however, this is not an appropriate measure. A measure such as mean squared error, or a non-parametric measure, might be appropriate.

**Fig S-3** evaluates the effects of volume conduction by implementing simulations to estimate the false positive rate as a factor of spatial resolution at the .265 threshold that is used by *Mendoza-Halliday et al*. for distant referencing, in comparison to bipolar and CSD derivations. Note that as spatial resolution decreases, false positive rate for the distant reference function increases rapidly. The same functions for bipolar and CSD derivations remain flat.

**Fig S-3.**
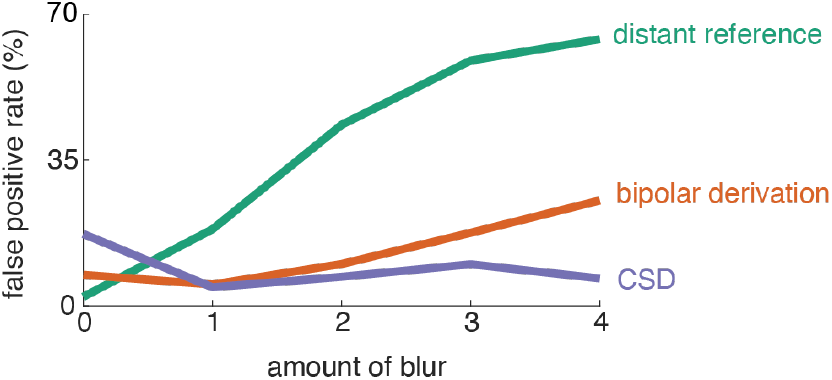
Increase in false positive rates for FLIP as a function of blur (volume conduction). Each point reflects 10,000 simulations of 30 channels with 1000 timepoints (simulating 1 second of data sampled at 1000 Hz) and 75 trials of 1/f pink noise (no patterns should be detected as these are random patterns) at each datapoint convolved with a 1-d blurring kernel to approximate volume conduction in the channel direction. For a distant reference, the false positive rate increases to over 60% with increasing blur. Bipolar referencing reduces the increase in false positives and Laplacian re-referencing keeps the false positive rate below 10% across these levels of blur.

## 4. Additional Broad Concerns

### 4.1. Conflation of oscillatory with sensory-evoked activity

*Mendoza-Halliday et al*. repeatedly attribute the reported motif to a laminar organization of alpha and gamma oscillations ^3,4^. Despite the focus on oscillatory activity, oscillations were not isolated from evoked responses. As described by *Mendoza-Halliday et al*., normalized relative power is calculated based on the -0.5 to 0.5 s interval around the onset of a stimulus. As such, the estimated relative power change does not distinguish between oscillations and evoked responses^5–8^. **Fig S-4A** depicts LFP spectrograms and power spectra in three channels in the supragranular, granular, and infragranular layer (2, 6, and 12, respectively) from *Mendoza-Halliday et al*.*’s* public dataset.

For the low frequencies, power was normalized to the mean in the -1 to 1 s interval, and for the high frequencies, power was normalized to the -1 to -0.25 s interval before stimulus onset. Evidently, the onset of the stimulus (indicated by the dashed line) is followed by a broad band high-frequency response in the gamma/BHA-band, as well as a low-frequency stimulus-evoked responses at 3 to 6 Hz. However, sustained ***<u>gamma-range oscillations</u>*** <u>are not observed in any of the channels</u>, which undermines claims about “oscillations,” and poses problems for subsequent work seeking to model the motif using gamma-specific mechanisms. The pre-stimulus period shows high power in the 15 to 20 Hz band, which is conventionally termed the beta-band. ***<u>Alpha oscillations</u>*** <u>(8 – 12 Hz) appear to be absent in the recording</u>. The spectrograms show a strong overlap between channels, indicating volume conduction due to the distant reference of the LFP. **Fig S-4B** depicts the Current Source Density (CSD) power spectra and demonstrates that the broadband response appears to be strongest in the supragranular layers, while the pre-stimulus beta activity is strongest in the granular layers. Stimulus-evoked and beta-band activity are attenuated in the infragranular channels.

***In sum***, the spectral analyses show that the spectrolaminar motif does not account for a contamination of oscillatory dynamics by evoked responses, and that the presence of ongoing oscillations in the alpha- and gamma-bands is not established. Numerous dedicated methods have been proposed that resolve oscillatory contributions to brain operations by: 1) identifying individual/local peak frequencies actually present in the data rather than blindly averaging over predefined bands^9^, 2) implementing tools to separate rhythmic and non-rhythmic contributions^10,11^, 3) eliminating spurious oscillatory activity due to harmonics and non-sinusoidal properties^12,13^, 4) characterizing transient burst events^14^, and 5) parametrizing cycle-by-cycle dynamics^15,16^. In general, traditional spectral methods used without such controls do not provide an accurate picture of underlying oscillatory dynamics^11^.

### 4.2. Conflation of <u>B</u>roadband <u>H</u>igh frequency <u>A</u>ctivity (BHA) with band-limited (30-60 Hz) gamma

Band-limited gamma oscillations (30 – 60 Hz) have been linked to coordinated interaction of pyramidal neurons and inhibitory interneurons^17,18^. Short bursts of activity in the higher 70-150 Hz range known as “high gamma” ^19,20^ or “broadband high frequency activity” (BHA) ^21,22^, contains a mixture of ripple (90-130 Hz) frequencies along with more wideband, transient (non-oscillatory) activity. BHA has further been shown to correlate strongly with multiunit activity ^19,21–23^, while band-limited gamma oscillations are associated with irregular firing patterns, also ^24^. Moreover, gamma oscillations and BHA have been shown to play distinct roles in perception and cognition ^25,26^. Due to the differences in the underlying mechanisms of gamma oscillations and BHA it is crucial to study these activities in isolation.

### 4.3. Inadequate consideration of the “State of the art” established by prior work

Contrary to statements in the Introduction^1^, at least 12 papers have analyzed laminar activity profiles across *multiple* cortical areas in monkeys ^5,27,36,37,28–35^ and in humans^38^. Interestingly, the Halgren paper^38^ proposes a cortical feedback role for alpha operating as a traveling wave through pyramidal neuron connections in the supragranular layers, which is a distinctly different mechanism from that advocated by *Mendoza-Halliday et al*.^1,3^. Bollimunta et al ^39^ utilize two terms that could be helpful here: (1) alpha generator, which refers to the active transmembrane current flow region, and (2) alpha pacemaker, which is the alpha generator that drives the other alpha generators. Though the generators appear in superficial (largest), middle and deep layers ^35^, the alpha pacemaker is in the deep layers in V2 and V4.

**Fig S-4.**
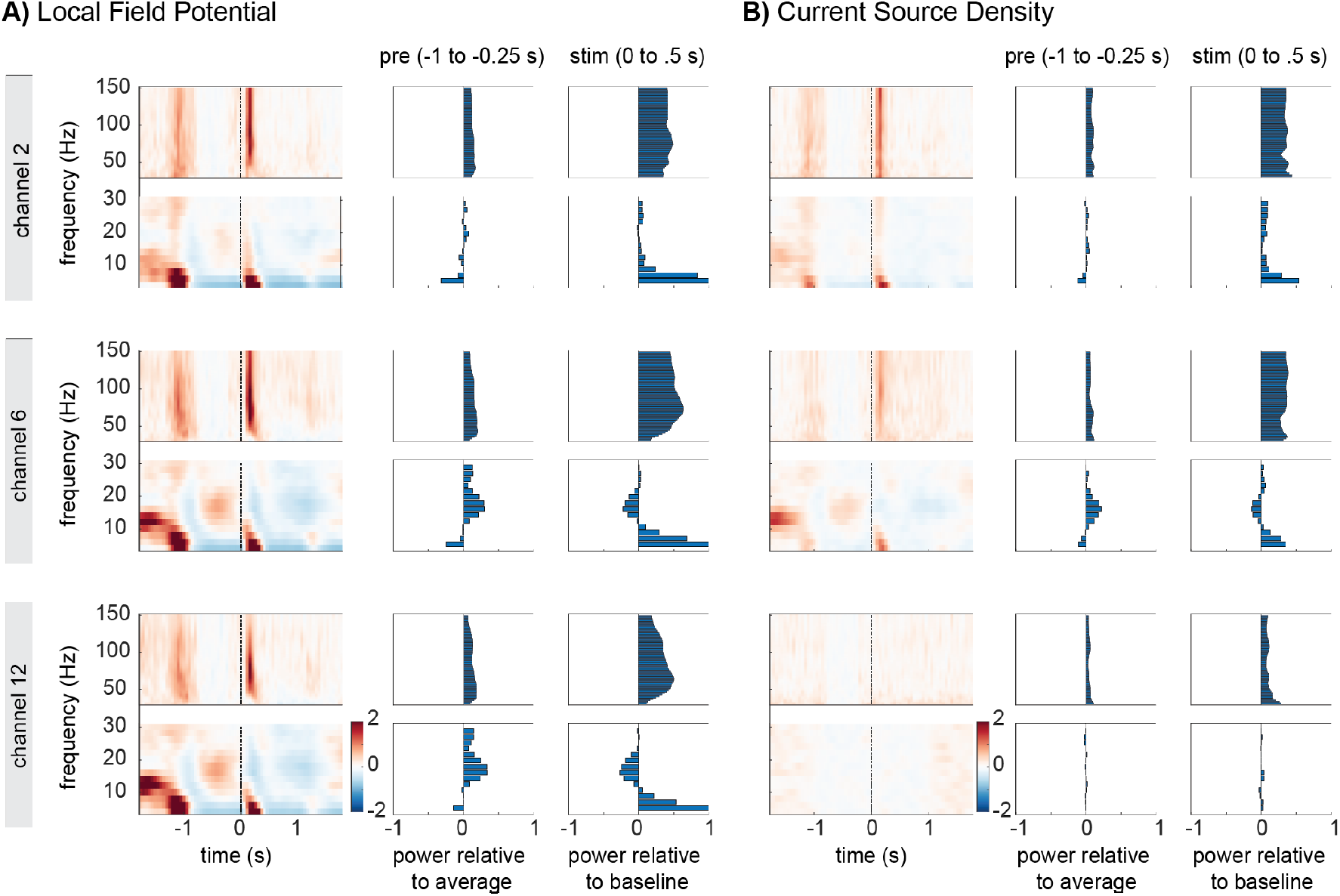
Oscillatory dynamics in the gamma- and alpha-bands are not established in *The Paper*. Spectrograms and spectra of the LFP, **A**) and Current Source Density (CSD, **B**) for three channels in the supragranular (2), granular (6), and infragranular layers (12), recorded from ventrolateral prefrontal cortex (vlPFC; publicly shared data from *The Paper*). Spectrograms are shown for the -1.75 to 1.75 s interval, with the stimulus onset at 0 seconds (dashed line). The spectra were calculated by averaging over the -1 to 0.25 s and 0 to 0.5 s intervals. For the low frequencies, power was normalized to the average power at each frequency in the channel, and for the high frequencies, power was normalized to the baseline intervals. **A)** The high-frequency spectrogram in the LFP demonstrates a broadband evoked response at 50 – 150 Hz following the onset of the stimulus. The evoked potential is indexed by the power increase in the 0 to 10 Hz band. The stimulus evoked activity is present in all trials, albeit strongest in the supragranular channel 2. The stimulus is preceded by high power in the beta-band (15-20 Hz) that is visible in all channels, but strongest in channel 6 and 12. **B)** CSD calculated from the LFP. Stimulus-evoked responses in both high and low-frequency ranges appear strongest in the supragranular channel 2. The pre-stimulus beta-band activity is strongest in the granular layer and disappears in the deep layers.

Another important methodological point evident from prior studies is that CSD profiles must be analysed on a case-by-case basis. Averaging across CSD profiles from different penetrations requires that the laminar profiles be perfectly aligned, which is rarely possible. Thus, averaging destroys information, produces circumscribed artifacts and yields physiologically implausible CSD profiles (see Fig 7 and Extended Fig 7). Findings from the earlier papers cited above, along with the large number of laminar recording studies focusing on single visual ^40–46^, somatosensory ^35,47–50^, auditory ^6,28,58–66,34,51–57^ and motor/prefrontal areas ^67–70^ collectively comprise a broader State-of-the-Art context in which to evaluate new ideas about cortex-wide motifs.

### 4.4. Ambiguous anatomical reconstructions

*Mendoza-Halliday et al*. provide single examples of histological sections from several cortical areas to illustrate methods by which key physiological findings (e.g., “gamma power peak” and laminar power function crossing points) are localized to specific layers in the cortex. Each example is followed by drawings that illustrate summary interpretations presumably based on viewing of the entire set of histological sections from all penetrations and lesions. Several concerns arise here.

***The broadest concern*** is that even acute penetrations by an electrode like those used here usually leave a prominent glial scar^29,44,71^, so that for a grid pattern of recordings, it is routine practice to reconstruct virtually all of the penetrations^34,72^. Such scars are not apparent at the sites that *Mendoza-Halliday et al*. represent as the electrode’s position (white circles mark probe channels) in their Fig. 4a,b and their Extended Data Fig.1. Possible electrode scars are sometimes visible (e.g., their Extended data Fig 4b) but not at the site and angle of the indicated probe location. The absence of a clear glial scar marking the electrode track suggests that the section used for illustration is not at the center of the electrolytical lesion, however, the reader cannot evaluate the quality, position and extent of lesions, as the critical adjacent sections are not provided.

#### Specific concerns arise from the histological illustrations

**1)** *<u>Lesions indicating the depth of the alpha/beta power peak in V1 and PMD are in the white matter</u>*. As *Mendoza-Halliday* note, this may be due to volume conduction from proximal areas. This localization deviates from the proposed motif, but it also underscores the risks (see **3**. above) inherent to the relatively poor spatial specificity of the LFP. **2)** *<u>Most of the five areas used for histological verification deviate in some way from the proposed motif</u>* (**Fig 4** and Extended data **Figs 3** and **4**). In LPFC, only 3-4/10 cases (30-40%) indicate power gradient crossings in L4. In PMD, the gamma power peak is depicted in L4 and the crossover is depicted in L5/6. In MST only 1 of the 2 cases provided depicts a power gradient crossing that is arguably in L4.

Overall, the histological presentation provides anecdotal and inconsistent, rather than quantitative and unequivocal evidence for laminar localization of critical physiological features. Given the emphasis *Mendoza-Halliday et al*. place on histological reconstruction as ground truth, a quantitative reconstruction of all sections showing evidence of lesions and electrode tracks would be required to alleviate this concern.

## 5. Physiologically realistic models to support laminar electrophysiology

To explain the inconsistencies between the laminar CSD and LFP profiles, the authors refer to previous computational work featuring a Neural Mass Model (NMM) that was developed based on a previously reported spectrolaminar motif in frontal cortex^3^. The NMM consists of two built-in alpha and gamma oscillators, constrained to produce band-limited dynamics at 10 and 40 Hz, respectively. However, neither the data presented^1^, nor in the referenced publication^3^ show band-limited peaks at 10 or 40 Hz (see above). While the NMM publication^3^ uses these band-limited frequencies to calculate the relative power gradient, FLIP and vFLIP select a wide range of frequencies to calculate the relative power for the slow and fast frequencies (see above). The model further relies on the assumption that the slow and fast oscillations are each generated by one single pyramidal population. However, *in vivo*, pyramidal populations exist across all layers and generate a wide range of frequencies^73,74^. Similarly, model synaptic input and return currents are located at the apical tuft and basal dendrites, whereas *in vivo* these currents can be spread throughout the dendritic tree^75^. Overall, the discrepancies between the NMM and the data indicate that the NMM may be ineffective at localizing the generators of the transmembrane currents responsible for the observed LFP/CSD patterns.

To accurately resolve laminar origins of LFP/CSD neuronal oscillations in an unbiased manner we propose instead constructing “informed” biophysical models utilizing detailed implementations of known cortical microcircuitry, multiple populations of excitatory (pyramidal IT, PT, CT, stellate) and inhibitory interneurons (SOM, PV, VIP, NGF, etc.), arrayed in realistic laminar cortical architecture, with precisely configured synaptic connection probabilities/locations, realistic dendritic morphologies, ion channel and synaptic time constants ^73,76^. Ideally also for dynamics like the alpha rhythm, models would include *thalamo*cortical circuitry^73,77^. These approaches create biophysical models that can simulate laminar LFPs/CSDs, using them to delineate population-level origin of neuronal oscillations^73,74,78^. Findings based on these approaches have demonstrated that LFP/CSD signals depend strongly on dendritic locations of synapses and show the importance of supragranular layers and inputs from other brain structures in creating alpha^74,79–81^. In the long run, these data-driven approaches will better complement empirical studies in resolving the laminar sources of neuronal dynamics including oscillations.

**Table S1.**
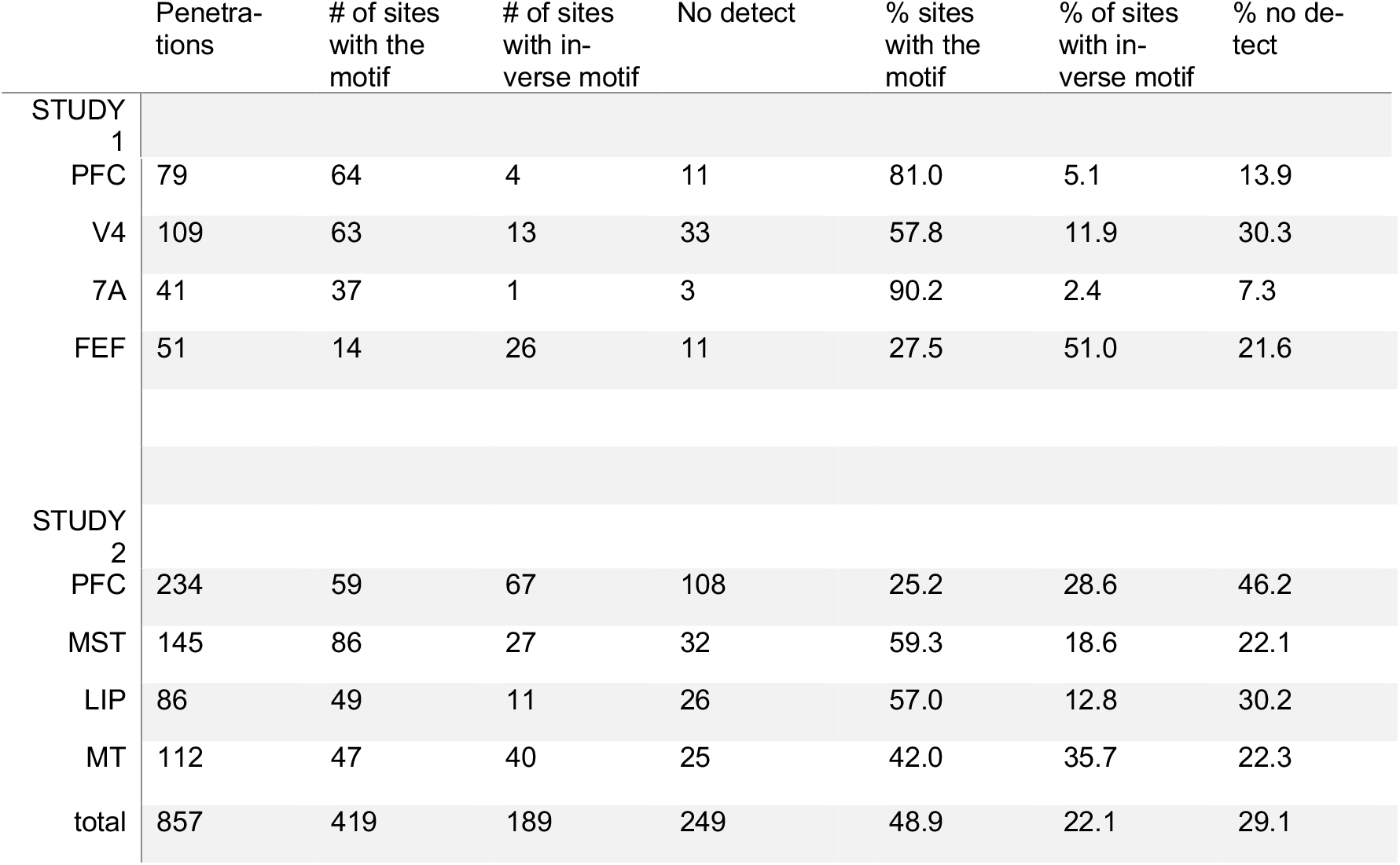
– significant FLIP fits from Mendoza-Halliday et al.’s dataset are not ubiquitous. Supplemental Table 1 – Summary of results from *The Paper*. The table indicates the number of penetrations detected by FLIP having the X-shaped “ubiquitous” motif for low and high frequency power across brain regions in the shared dataset from *The Paper*. Substantial interregional and inter-study variability can be observed, such as only 25% of PFC sites exhibiting the motif in Study 2.

|  | Penetra-<br>tions | # of sites<br>with the<br>motif | # of sites<br>with in-<br>verse motif | No detect | % sites<br>with the<br>motif | % of sites<br>with in-<br>verse motif | % no de-<br>tect |
| --- | --- | --- | --- | --- | --- | --- | --- |
| STUDY<br>1 |  |  |  |  |  |  |  |
| PFC | 79 | 64 | 4 | 11 | 81.0 | 5.1 | 13.9 |
| V4 | 109 | 63 | 13 | 33 | 57.8 | 11.9 | 30.3 |
| 7A | 41 | 37 | 1 | 3 | 90.2 | 2.4 | 7.3 |
| FEF | 51 | 14 | 26 | 11 | 27.5 | 51.0 | 21.6 |
| STUDY 2 |  |  |  |  |  |  |  |
| PFC | 234 | 59 | 67 | 108 | 25.2 | 28.6 | 46.2 |
| MST | 145 | 86 | 27 | 32 | 59.3 | 18.6 | 22.1 |
| LIP | 86 | 49 | 11 | 26 | 57.0 | 12.8 | 30.2 |
| MT | 112 | 47 | 40 | 25 | 42.0 | 35.7 | 22.3 |
| total | 857 | 419 | 189 | 249 | 48.9 | 22.1 | 29.1 |

